# Causality-Enriched Epigenetic Age Uncouples Damage and Adaptation

**DOI:** 10.1101/2022.10.07.511382

**Authors:** Kejun Ying, Hanna Liu, Andrei E. Tarkhov, Marie C. Sadler, Ake T. Lu, Mahdi Moqri, Steve Horvath, Zoltán Kutalik, Xia Shen, Vadim N. Gladyshev

## Abstract

Machine learning models based on DNA methylation data can predict biological age but often lack causal insights. By harnessing large-scale genetic data through epigenome-wide Mendelian Randomization, we identified CpG sites potentially causal for aging-related traits. Neither the existing epigenetic clocks nor age-related differential DNA methylation are enriched in these sites. These CpGs include sites that contribute to aging and protect against it, yet their combined contribution negatively affects age-related traits. We established a novel framework to introduce causal information into epigenetic clocks, resulting in DamAge and AdaptAge—clocks that track detrimental and adaptive methylation changes, respectively. DamAge correlates with adverse outcomes, including mortality, while AdaptAge is associated with beneficial adaptations. These causality-enriched clocks exhibit sensitivity to short-term interventions. Our findings provide a detailed landscape of CpG sites with putative causal links to lifespan and healthspan, facilitating the development of aging biomarkers, assessing interventions, and studying reversibility of age-associated changes.

## Introduction

Aging, a complex biological process, is characterized by the accumulation of deleterious molecular changes, leading to organ and system function decline and eventual death ^1^. Although the underlying mechanisms of aging are not well understood, various studies indicate that aging is associated with altered DNA methylation patterns, particularly 5-methylcytosine (5mC) in mammals, where overall methylation slightly decreases in adulthood, but certain regions may experience hypo- or hypermethylation ^2,3^. Specific CpG site methylation levels correlate strongly with age, enabling the development of machine learning-based epigenetic aging clocks (e.g., Horvath pan tissue and Hannum blood-based clocks) that predict biological sample ages with high accuracy ^4–6^. These clocks, more closely related to health measures than chronological age, are seen as better biological age indicators ^7^.

However, epigenetic aging clocks, primarily correlative, should be interpreted cautiously ^8^. It remains unclear whether the DNA methylation differences used for age prediction are causal to aging-related phenotypes or merely aging byproducts. Establishing causality typically involves randomized controlled trials (RCTs), where treatment effects are assessed by comparing intervention and control groups, eliminating confounding factors ^9,10^. But, given the genome’s vast CpG sites, such individual perturbations are impractical.

Mendelian randomization (MR), a genetic causal inference approach, mimics RCT principles ^11^. It uses genetic variants associated with exposure as instrumental variables, leveraging the natural random distribution of parental genome to offspring. MR has been applied to molecular traits using molecular quantitative trait loci (molQTLs), including DNA methylation (meQTL) ^12^. Previous studies have successfully used meQTLs to identify causally linked CpG sites for diseases ^13^. Integrating molQTLs with genome-wide association studies for aging-related traits allows for two-sample MR to estimate the molecular changes’ causal effects on aging.

Here, we leveraged large-scale genetic data and performed epigenome-wide Mendelian Randomization (EWMR) on 420,509 CpG sites to identify CpG sites that are causal to eight aging-related traits. We found that none of the existing clocks are enriched for putative causal CpG sites. We further constructed a causality-enriched clock based on this inferred causal knowledge, as well as clocks that separately measure damaging and protective changes. Their applications provide direct insights into the aging process. Thus, our results offer a comprehensive map of human CpG sites causal to aging traits, which can be used to build causal biomarkers of aging and assess novel anti-aging interventions and aging-accelerating events.

### Epigenome-wide MR on aging-related phenotypes

MR is an established genetic approach for causal inference that utilizes natural genetic variants as instrument variables ^14^. Since the allocation of genetic variants is a random process and is determined during conception, the causal effects estimated using MR are not biased by environmental confounders. Therefore, it could be used as a tool for investigating causal relationships between the DNA methylation and aging-related phenotypes (Fig. 1a). In the context of MR, a CpG site can be defined as causal when associations of SNPs with CpG methylation are directionally consistent and proportional in magnitude to its associations with aging-related phenotypes.

**Fig. 1.**
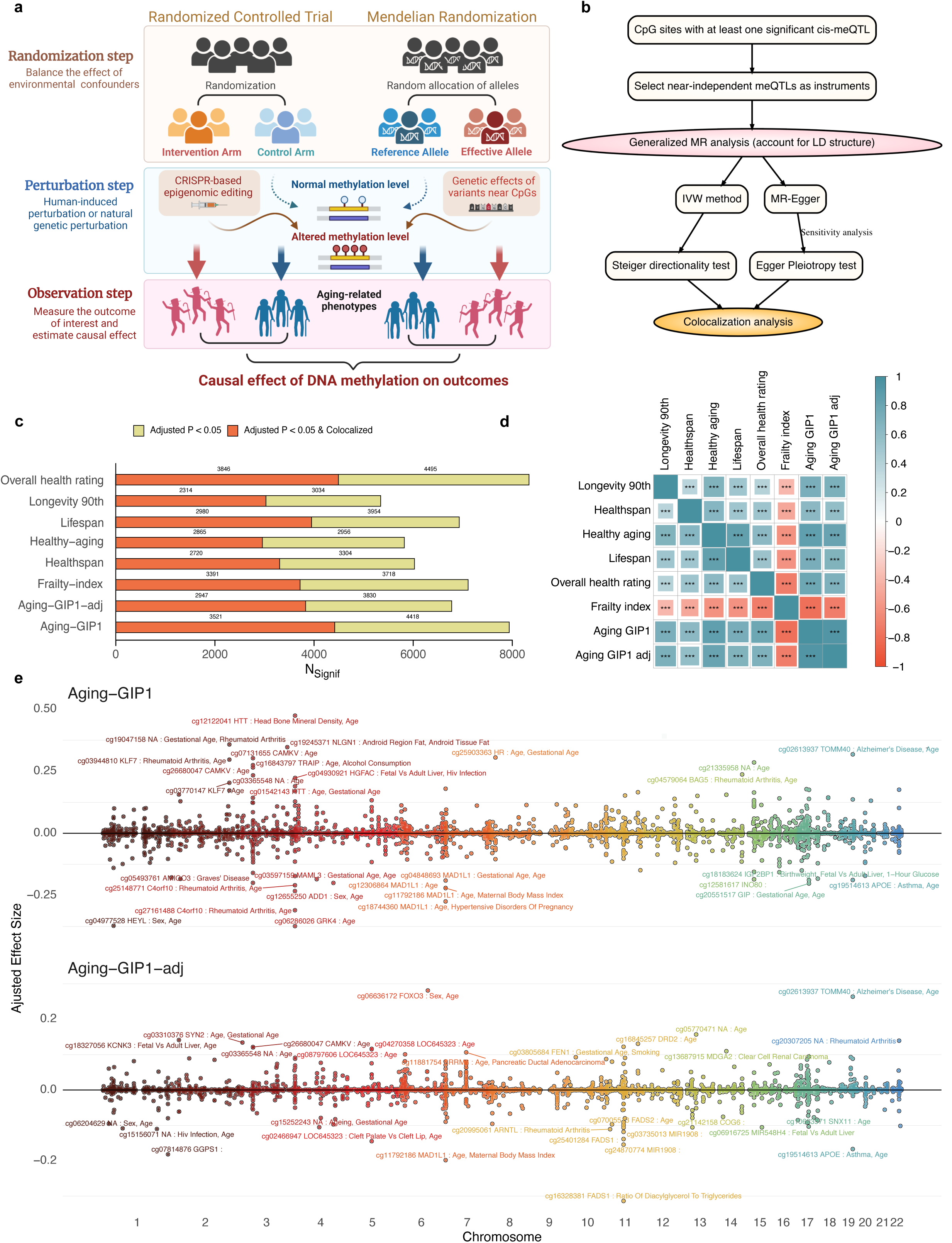
Epigenome-wide Mendelian Randomization on various aging-related phenotypes. **a.** Schematic diagram shows the principle of MR using meQTLs as exposures and aging-related traits as outcomes to identify putative causal CpG sites. **b.** Flow chart shows the procedure for epigenome-wide MR and sensitivity analysis. **c.** Number of significant putative causal CpG sites identified for each trait after adjusting for multiple comparisons using the Bonferroni correction. Red regions of the bars indicate the number of putative causal CpG sites supported by the colocalization analysis with conditional PP-H4 > 0.7. **d.** Spearman correlation of the estimated causal effects of CpGs in twelve traits. *P* values reported are adjusted for multiple comparisons using Bonferroni correction. Only CpGs with significant MR signals across at least six traits are included in the analysis. Color scheme reflects Spearman correlation coefficients, * adjusted *P* < 0.05, ** adjusted *P* < 0.01, *** adjusted *P* < 0.001. **e.** Modified Mississippi plot shows significant MR signals for Aging-GIP1 and adjusted Aging-GIP1. Only the CpG sites adjusted for multiple comparisons with FDR < 0.05 are shown in the plot. X-axis corresponds to the genomic positions of CpG sites; Y-axis represents the size of the causal effect adjusted by colocalization probability (PP-H4). CpG sites with top adjusted causal effects are annotated with the name and nearest gene. Only CpG sites with adjusted *P* < 0.05 are included in the plot.

To identify CpG sites causal to aging, we used 420,509 CpG sites with meQTLs available (GoDMC, whole blood samples from 36 cohorts, 27,750 European subjects) as exposures and selected eight aging-related phenotypes as outcomes (Fig. 1a, Methods, Table 1), including two lifespan-related traits: lifespan and extreme longevity (defined as survival above 90th percentile) ^15,16^; three health-related traits: healthspan (age at the first incidence of any major age-related disease) ^17^, frailty index (measurement of frailty based on the accumulation of a number of health deficits during the life course) ^18^, and self-rated health (based on the questionnaire responses) ; and three summary-level aging-related traits: Aging-GIP1 (the first genetic principle component of six human aging traits - healthspan, father’s and mother’s lifespan, exceptional longevity, frailty index and self-rated health) ^19^, socioeconomic traits-adjusted Aging-GIP1, and healthy aging (multivariate genomic scan of healthspan, lifespan, and longevity) ^20^.

**Table 1.** Datasets used in this study.

| Dataset | Description |
| --- | --- |
| *GWAS data* |  |
| meQTLs | meQTLs were obtained from the Genetics of DNA Methylation Consortium (GoDMC). DNA methylation levels were measured in whole blood samples from 36 cohorts, including 27,750 European subjects. 420,509 CpG sites were analyzed (Min et al.)<sup>12</sup>. |
| Aging-GIP1 | First genetic principal component of six human aging traits - healthspan, father and mother lifespan, exceptional longevity, frailty index and self-rated health, which captures both length of life and indices of mental and physical wellbeing (Timmers et al.)<sup>19</sup>. |
| Aging-GIP1-adj | Aging-GIP1 adjusted for household income and socioeconomic deprivation, from the same GWAS study as above<sup>19</sup>. |
| Healthy aging | The multivariate genomic scan of healthspan, lifespan, and longevity (Timmers et al.)<sup>20</sup>. |
| Lifespan | GWAS of lifespan from 512,047 mothers and 500,193 fathers of European ancestry (Timmers et al.)<sup>15</sup>. |
| Longevity | 11,262 subjects of European ancestry with a lifespan above the 90th percentile as the case group and 25,483 control subjects whose age at the last visit was below the 60th percentile age (Deelen et al.)<sup>16</sup>. |
| Healthspan | The age of the first incidence of any major age-related disease, including dementia, congestive heart failure, diabetes, chronic obstructive pulmonary disease, stroke, cancer, myocardial infarction, as well as the incidence of death. The GWAS of healthspan included 300,447 subjects of European ancestry from the UK Biobank cohort (Zenin et al.)<sup>17</sup>. |
| Frailty index | Calculated based on the cumulative number of health deficits during aging. The frailty index GWAS included 164,610 UK Biobank participants aged 60-70 years and 10,616 Swedish TwinGene participants aged 41-87 years (Atkins et al.)<sup>18</sup>. |
| Self-rated health | Self-rated health GWAS was based on questionnaire responses on a scale of 0-5 in UK Biobank cohort, downloaded from Pan-UKBB project. |
| *GEO data* |  |
| GSE107143 | This study conducted DNA methylation analyses of blood samples from atherosclerosis patients and healthy donors. |
| GSE127985 | DNA methylation changes in prostate cancer cases and it's prognosis. |
| GSE192918 | This study analyzed peripheral whole blood DNA methylation profiles of pregnant women at different stages of gestation and post-delivery, identifying changes in DNA methylation patterns associated with different time points during pregnancy. |
| GSE193795 | Genome-wide DNA methylation profiling was performed on 44 hypertensive and 44 healthy control samples, revealing distinct DNA methylation patterns associated with hypertension. |
| GSE210245 | DNA methylation data from human whole blood samples were analyzed to assess the impact of treatment with human umbilical cord plasma concentrate injection. |
| GSE51954 | Genome-wide DNA methylation profiling was conducted on epidermal and dermal samples obtained from sun-exposed and sun-protected body sites. |
| GSE94876 | This study compared global methylation changes in buccal cells between smokers, moist snuff consumers, and non-tobacco consumers. |
| GSE98056 | This study aimed to explore genome-wide DNA methylation changes and identify altered biological pathways resulting from n-3 fatty acid supplementation in overweight and obese individuals. |
| GSE101673 | DNA methylation data for cigarette smoke condensate treated cell. |
| GSE78773 | This study identified multi-locus methylation disturbances in individuals with different methylation patterns, including patients with Temple and Angelman syndromes |
| GSE90117 | This study identified the relationship between PON1 activity, allele, and DNA methylation. |
| GSE79257 | This study analyzed DNA methylation in infants born through different assisted reproductive techniques and unassisted conception, utilizing archived Guthrie cards for methylation profiling. |
| GSE42865 | This study analyzed DNA methylation in B cells from patients with Hutchinson-Gilford Progeria Syndrome (HGP) and Werner Syndrome and controls. |

Aging-GIP1 captures both the length of life and age-related health status, which can be considered as a genetic representation of healthy longevity. It also shows the strongest genetic correlation with all other traits related to lifespan ^19^. Therefore, we further used Aging-GIP1 as the primary aging-related trait to investigate CpG sites causal to the aging process. A genetic correlation analysis showed that all eight lifespan- and health-related traits are genetically correlated and clustered with each other (Extended Data Fig. 1).

We then applied generalized inverse-variance weighted MR (gIVW) and MR-Egger (gEgger) to each exposure-outcome pair (Method). Post-Bonferroni correction, we identified over 6,000 significant causal CpG sites for each trait (Fig. 1c). The causal effect estimates across traits exhibited a strong correlation (Fig. 1d). To mitigate reverse causation, we conducted cis-reverse MR and Steiger directionality tests ^21^. Additionally, to control false positives from MR due to linkage disequilibrium (LD) or pleiotropy, we also employed pairwise conditional and colocalization (PWCoCo) analysis (Fig. 1b, Method) ^22^. Over half of the MR-identified CpG sites for each trait showed colocalized signals.

Given the analysis was limited to 420,509 CpG sites, the effects of unmeasured CpG sites on traits couldn’t be differentiated. To validate whether MR-estimated effects can be attributed to single CpG sites, we analyzed naturally occurring point mutations at putative causal CpG sites (meSNP). Nearly 10% of CpG sites on the human methylation array have an meSNP, and we found significant depletion of meSNPs at causal sites, suggesting negative selection against loss-of-function mutations, likely due to causal site enrichment in regulatory regions (Extended Data Fig. 2). A significant positive correlation (Pearson’s R = 0.4, P = 1e-4) was observed between outcomes estimated using single meSNPs and MR for causal CpG sites with available meSNPs (Extended Data Fig. 2). Thus, the MR-estimated causal effects could partially be ascribed to individual CpG sites, yet many without meSNPs, and their methylation levels being highly correlated with neighboring sites, suggest these sites tag causal regulatory regions.

To prioritize CpG sites for Aging-GIP1, we filtered MR signals using P value thresholds after Bonferroni correction and ranked them by causal effect magnitude and colocalization probability (PP.H4). Top pro-longevity CpG sites included cg12122041 (HTT) annd cg02613937 (TOMM40) while top sites limiting longevity were cg04977528 (HEY) and cg06286026 (GRK4). Additionally, cg19514613 at the APOE locus was significant for limiting longevity. Notably, lifespan effects at HTT and MAML3 loci were previously identified ^19^, and TOMM40 and APOE are linked to Alzheimer’s disease and lifespan. Adjusting for socioeconomic status, cg06636172 at the FOXO locus emerged as the top pro-longevity CpG site (Fig. 1e).

### Putative causal CpGs are enriched in regulatory regions

To further understand properties of the CpG sites identified as causal to each aging-related trait, we performed an enrichment analysis using 14 Roadmap annotations ^23^. We found that the putative causal CpGs for most traits are enriched in promoters and enhancers while depleted in quiescent regions (Fig. 2a). Furthermore, these sites were enriched in CpG shores (Extended Data Fig. 3). We observed that the putative causal CpG sites for Aging-GIP1 are significantly more evolutionally conserved compared to non-causal CpGs, based on both functional genomic conservation scores (Learning Evidence of Conservation from Integrated Functional genomic annotations, LECIF) and the phastCons/phyloP scores across 100 vertebrate genomes ^24^ (Fig. 2b, c, Extended Data Fig. 4). Moreover, the absolute value of the estimated causal effect sizes showed significant positive correlations between all three conservative scores. These results suggest that the CpG sites identified as causal for aging-related traits are more likely to be in functional genomic elements and more evolutionarily conserved.

**Fig. 2.**
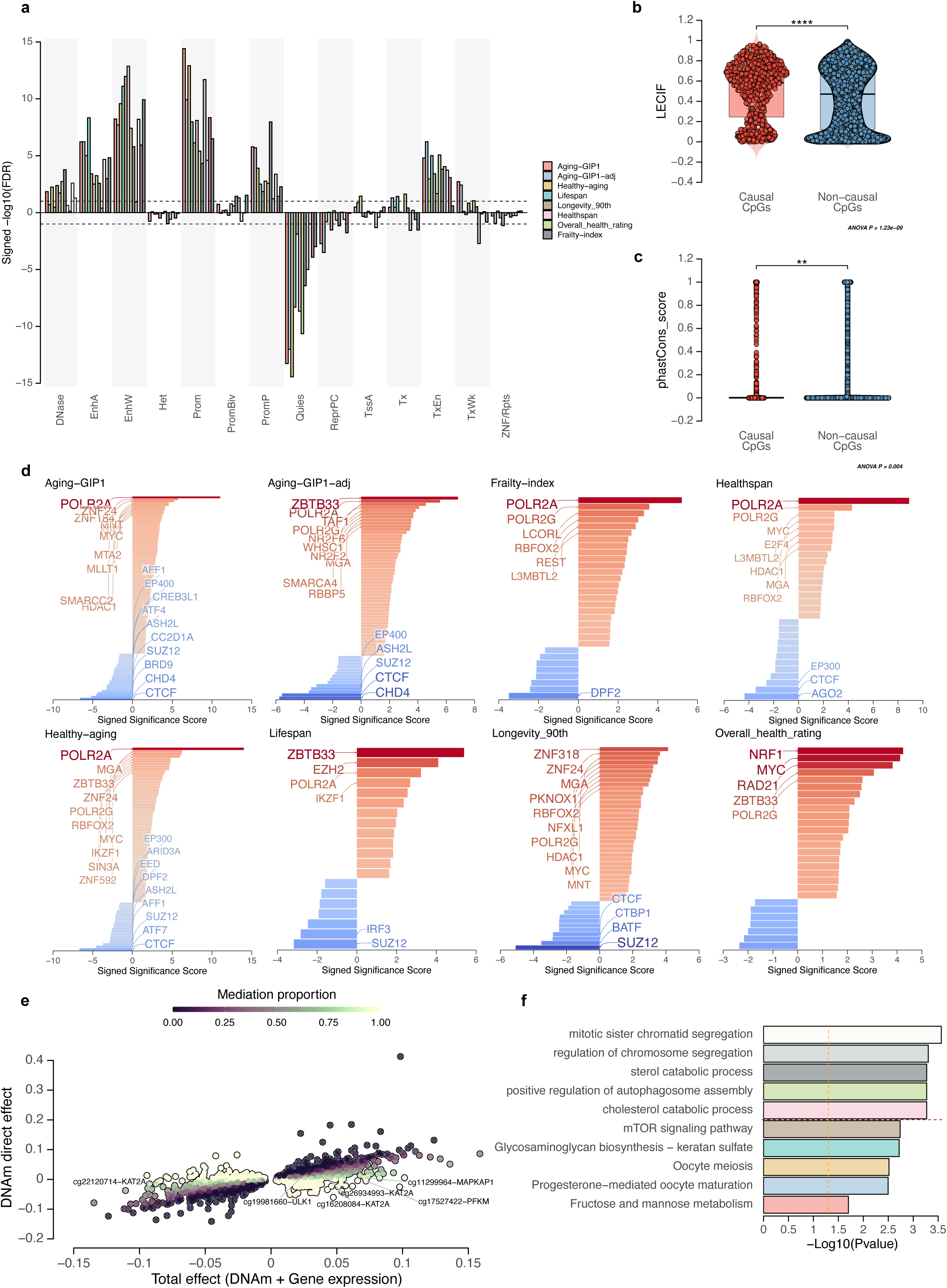
CpG sites causal to aging are enriched in specific genetic regulatory regions. **a.** Bar plot shows enrichment of putative causal CpG sites in 14 Roadmap genomic annotations. Y axis shows -log10 (FDR) based on Fisher’s exact test, signed by log2 (Odds ratio). Putative causal CpG sites identified for different traits are annotated with different colors. Two dotted horizontal lines show the FDR threshold of 0.05. TssA, active transcription start site. Prom, upstream/downstream TSS promoter. Tx, actively transcribed state. TxWk, weak transcription. TxEn, transcribed and regulatory Prom/Enh. EnhA, active enhancer. EnhW, weak enhancer. DNase, primary DNase. ZNF/Rpts, state associated with zinc finger protein genes. Het, constitutive heterochromatin. PromP, Poised promoter. PromBiv, bivalent regulatory states. ReprPC, repressed polycomb states. Quies, quiescent state. **b, c.** Box plot shows distribution of conservation scores in causal (n = 1053) and non-causal CpG sites (n = 161881) for Aging-GIP1. Two-sided t-test was used. Conservation scores are obtained by Learning Evidence of Conservation from Integrated Functional genomic annotations (LECIF, **b**) and phastCons (**c**). * *P* < 0.05, ** *P* < 0.01, *** *P* < 0.001, **** *P* < 0.0001. **d.** Enrichment of putative causal CpG sites for 12 aging-related traits against transcription-factor-binding sites. Each horizontal bar represents an enriched term. The X-axis shows the - log10(*P*-value), signed by log2 (Odds ratio). The top 10 enriched terms that passed the FDR threshold of 0.05 for each direction are annotated. P values are adjusted for multiple testing using FDR method. **e.** Scatter plot showing the mediation analysis of Aging-GIP1. The total causal effects are shown on X-axis and the direct effects of DNA methylation are shown on Y-axis. The color shows the proportion of DNAm causal effect that is mediated by gene expression. Top CpG-gene pairs are annotated. **f.** Enrichment of the top mediator gene for Aging-GIP1 in GO terms (above dashed line) and KEGG pathways (below the dashed line). The X-axis shows -log(P) for the fisher exact test. One-sided test was used and the P-values are adjusted for multiple testing using FDR method.

It is well known that DNA methylation status may affect transcription factor (TF) binding. To understand the relationship between putative causal CpG sites and TFs, we performed a transcription factor binding site enrichment analysis (Fig. 2d). The CpG sites causal to Aging-GIP1 were significantly enriched in the binding sites of 63 TFs, including POLR2A, ZNF24, MYC, and HDAC1; while depleted in the binding sites of 19 TFs, including CTCF, CHD4, and BRD9 (Fig. 2d). In particular, POLR2A was among the top enriched TFs in 6 of 8 traits. POLR2A is the POLR2 subunit (RNA polymerase II), and previous research showed that epigenetic modifications can modulate its elongation and affect alternative splicing ^25^. Our results imply that this mechanism is potentially a major contributor that mediates the effects of DNA methylation on aging. We further found that there were 3 TF-binding sites (BRD4, CREB1, and E2F1) enriched with CpG sites whose methylation levels promote healthy longevity (Aging-GIP1), and 4 TF-binding sites (HDAC1, ZHX1, IKZF2, and IRF1) enriched with CpG sites whose methylation levels decrease healthy longevity (Extended Data Fig. 4).

Since the putative causal CpGs are enriched in regulatory regions and TF binding sites, we further performed a mediation analysis to investigate whether the effect top CpG hits are mediated through gene expression. The mediation effects were estimated through multivariable MR including both DNA methylation and gene expression, which dissect significant CpG-phenotype causal effects (θ_T_) into direct effects (θ_D_) and indirect effects through transcript levels (Method) ^26^. Among 2,255 putative causal CpGs applicable to the mediation analysis for Aging-GIP1, 1,000 had their effect mediated by a major transcript (with mediation proportion > 0.03, Fig. 2e). For example, we found that the 92% of the effect of cg11299964 on Aging-GIP1 is mediated through the expression of MAPKAP1, which is a key protein in the mTOR signaling pathway (Fig. 2e). We then performed a gene set enrichment analysis on GO and KEGG using the mediator genes for Aging-GIP1 (Fig. 2f). We found that the mediators are enriched in several aging-related pathways, including mTOR signaling (P = 0.0018) and autophagosome assembly (P = 5.4e-4, Fig. 2f).

We also examined the enrichment of putative causal CpG sites in phenome-wide EWAS signals obtained from the EWAS catalog ^27^. The top enriched phenotypes included rheumatoid arthritis, HIV infection, nitrogen dioxide exposure, and maternal obesity (Extended Data Fig. 5). Interestingly, none of these conditions is primarily caused by aging but instead increased risk of mortality. Therefore, our results suggest that the causal CpG sites for aging are enriched in conditions that cause accelerated aging, but not in conditions that are caused by aging, consistent with the previous study that suggests that differentially expressed genes reflect disease-induced rather than disease-causing changes ^1^.

### Existing Epigenetic clocks lack aging-causal CpGs

One open question for epigenetic clocks is whether their clock sites are causal to aging and age-related functional decline. To answer this question, we collected seven epigenetic age models in humans, namely, the Zhang clock, PhenoAge, GrimAge, PedBE, HorvathAge, HannumAge, and DunedinPACE. We then performed an enrichment analysis of putative causal CpGs for all eight lifespan/healthspan-related traits for each clock. After correcting for multiple testing, none of the existing clocks showed significant enrichment for putative causal CpGs of any of the lifespan/healthspan-related traits (Fig. 3g). PhenoAge showed a nominal significant enrichment with CpGs causal to healthspan and healthy aging, but it was not robust to the choice of thresholds. This finding suggests that, although some clocks contain CpGs causal to aging (Table 2), they, by design, favor CpG sites with a higher correlation with age and thus are not enriched with putative causal CpGs.

**Fig. 3.**
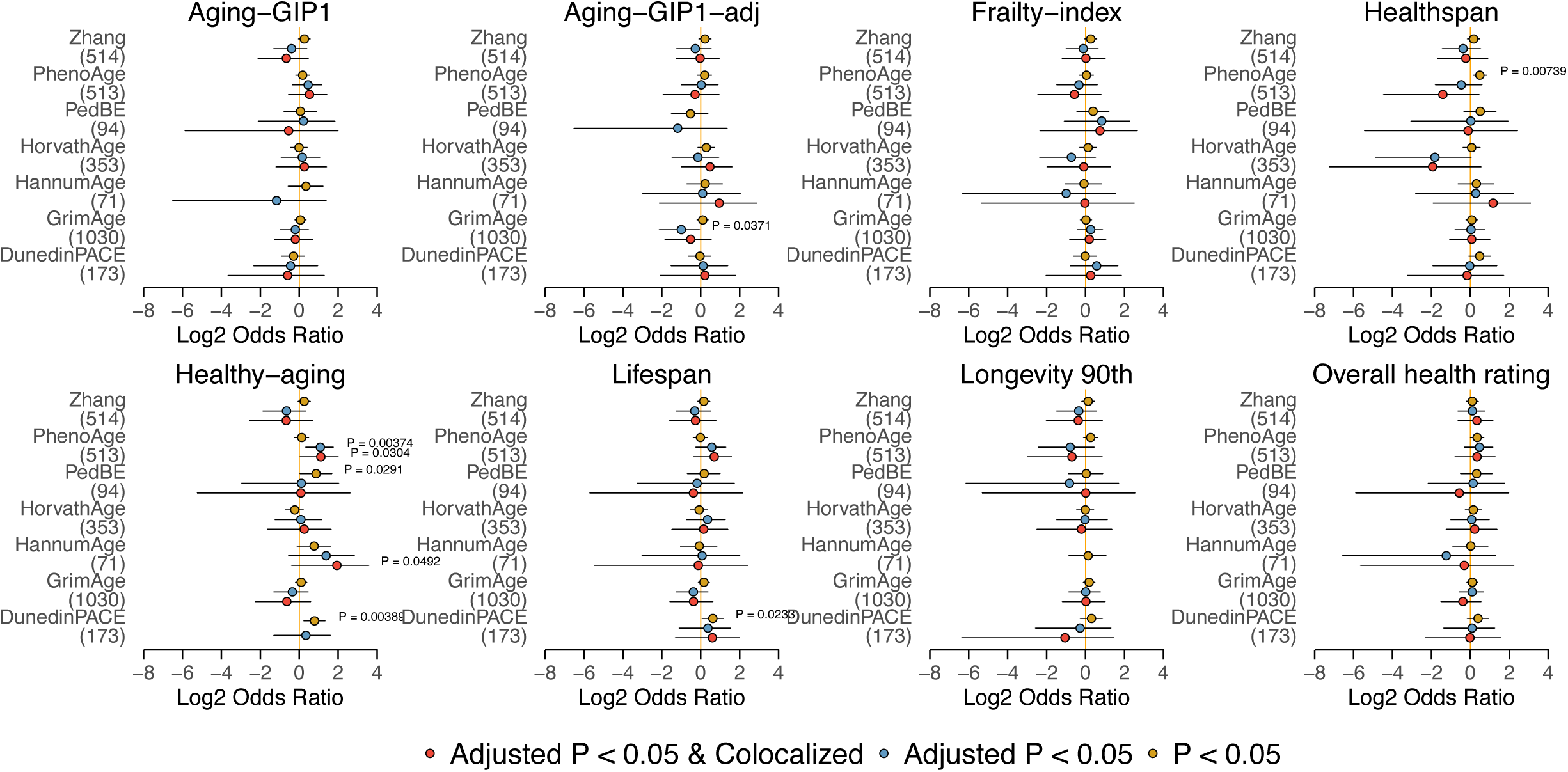
Existing epigenetic clocks are not enriched with CpG sites causal to aging. Forest plot shows enrichment of clock sites for six aging clock models in putative causal CpG sites identified by MR for each trait. N of CpG sites for each clock model are annotated in parenthsis under the name of each clock. X-axis shows the log2(Odds Ratio). *P*-values calculated by Fisher’s exact test are annotated if *P* < 0.05. The dot shows the estimated value of log2(Odds Ratio). Error bars show 95% confidence intervals. Different colors represent the different thresholds for putative causal CpGs. Two-sided Fisher exact test was used. Raw P-values are annotated as none of the results passed the FDR threshold of 0.05.

**Table 2.** Putative causal CpG sites in existing epigenetic clocks. P values are adjusted for multiple testing using Bonferroni’s method.

|  | Position | Weight | outcome | Beta | SE | P | H4 | role |
| --- | --- | --- | --- | --- | --- | --- | --- | --- |
| HorvathAge (353) | cg06557358 | -0.14 | Overall_health_rating | -0.04 | 0.008 | 1.96E-07 | 0.89 | Protective |
|  | cg09509673 | 0.01 | Healthy-aging | 0.02 | 0.003 | 3.86E-13 | 0.85 | Protective |
|  | cg09509673 | 0.01 | Lifespan | 0.05 | 0.006 | 9.92E-20 | 0.83 | Protective |
|  | cg11299964 | -0.16 | Aging-GIP1 | 0.08 | 0.012 | 5.42E-12 | 0.86 | Deleterious |
|  | cg16744741 | -0.35 | Aging-GIP1 | 0.09 | 0.017 | 1.86E-08 | 0.89 | Deleterious |
|  | cg16744741 | -0.35 | Overall_health_rating | 0.06 | 0.008 | 6.10E-14 | 0.86 | Deleterious |
| PhenoAge (513) | cg05087948 | -6.99 | Aging-GIP1-adj | -0.08 | 0.013 | 7.30E-10 | 1.00 | Protective |
|  | cg21926612 | -2.15 | Overall_health_rating | 0.01 | 0.002 | 3.27E-11 | 0.94 | Deleterious |
|  | cg11896923 | -1.38 | Healthspan | 0.17 | 0.024 | 4.64E-12 | 0.90 | Deleterious |
|  | cg11896923 | -1.38 | Healthy-aging | 0.05 | 0.008 | 5.63E-10 | 0.86 | Deleterious |
|  | cg00862290 | -0.23 | Healthy-aging | 0.00 | 0.001 | 1.28E-08 | 0.85 | Protective |
|  | cg00862290 | -0.23 | Lifespan | -0.02 | 0.003 | 0 | 0.94 | Protective |
| Zhang (514) | cg24987259 | -1.33 | Overall_health_rating | -0.04 | 0.007 | 8.25E-09 | 0.95 | Protective |
|  | cg05310309 | 0.18 | Aging-GIP1 | 0.03 | 0.003 | 1.13E-32 | 0.96 | Protective |
|  | cg05310309 | 0.18 | Overall_health_rating | 0.01 | 0.002 | 2.64E-12 | 0.92 | Protective |
|  | cg06672696 | 0.02 | Frailty-index | 0.05 | 0.010 | 1.74E-07 | 0.82 | Protective |
| PedBE (94) | cg04221461 | 0.03 | Frailty-index | 0.04 | 0.008 | 1.25E-07 | 0.95 | Protective |
|  | cg19381811 | -0.08 | Aging-GIP1 | -0.04 | 0.004 | 3.26E-21 | 0.929544 | Protective |
|  | cg19381811 | -0.08 | Overall_health_rating | -0.03 | 0.002 | 8.80E-37 | 0.955032 | Protective |

### Identify protective and deleterious DNAm changes in aging

Another important question in epigenetic aging is the identity and number of epigenetic changes that (i) contribute to age-related damage and (ii) respond to it ^30^. We approached this question by integrating information on the causal effect and age-related differential methylation for each CpG. The protective or damaging nature of age-related differential methylation at each CpG is indicated by the product of the causal effect and age-associated differential methylation (*b*_*age*_ × *b*_*MR*_, Fig. 4a). For example, if a higher methylation level of a certain CpG site leads to a longer lifespan or healthspan, then during aging, a decrease of the methylation level at that site would be considered as having a damaging effect, whereas an increased methylation level would be considered as having a protective effect.

**Fig. 4.**
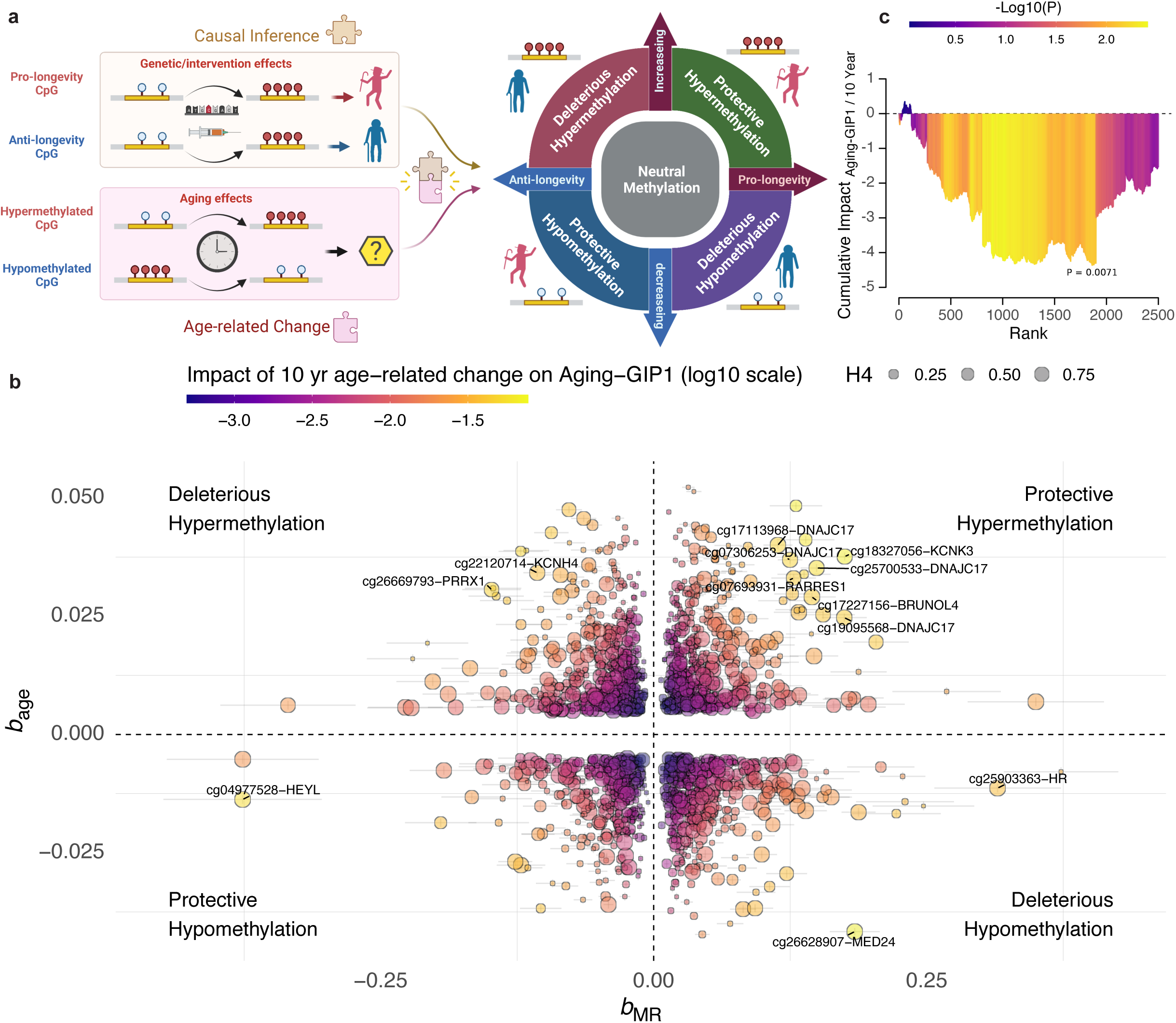
Integration of causal information and age-associated differential methylation to separate protective and damaging epigenetic changes. **a.** Schematic diagram showing the method to identify protective and damaging epigenetic changes by integrating MR results and age-related differential methylation. b. Relationship between MR-estimated causal effects (X-axis) and age-related differential methylation (Y-axis) for each significant putative causal CpG identified in Aging-GIP1. The color scheme highlights the expected impact of age-related differential methylation on aging. Error bars show the standard error of *b*. The size reflects the PP-H4. Only CpG sites with adjusted *P*-values < 0.05 and relative PP-H4 > 0.7 are plotted. The CpG sites with the top 10 largest effect sizes are annotated. **c.** Area plots show the total cumulative effect of changes in DNA methylation on Aging-GIP1. X-axis shows the rank of top 3,000 CpG sites based on the magnitude of age-associated differential methylation. Y-axis and the color scheme show the P-value estimated by 10,000 permutation tests. Two-sided Fisher exact test was used.

The effect of DNA methylation estimated by MR is estimated through linear regression, which assumes that the relationship between DNA methylation level and lifespan-related outcome is linear. Prior to annotating protective and damaging CpGs, it is important to make sure the effect size of genetic instruments on DNA methylation levels is in the same order of magnitude as the effect of aging. We show that the effect of genetic instruments is comparable with the effect of aging by calculating the ratio between the effect of strongest cis-meQTL and age-related differential methylation for each CpG site (Extended Data Fig. 6). The median ratio is 21.8 for all significant age-associated sites and 3.9 for top 50 age-associated sites, suggesting that the median effect of genetic instruments is roughly equivalent to the effect of years of aging.

Therefore, using the age-related blood DNA methylation data estimated from 7,036 individuals (ages of 18 and 93 years, Generation Scotland cohort)^31^, we separated the CpG sites causal to eight traits related to lifespan into four different categories: protective hypermethylation, deleterious hypermethylation, protective hypomethylation, and deleterious hypomethylation (Fig. 4b, Extended Data Fig. 7). Among the top 10 CpG sites whose differential methylation during aging has a relatively large impact on healthy longevity, six hypermethylated CpG sites during aging exhibit strong protective effects, including cg18327056, cg25700533, cg19095568, cg17227156, cg17113968, and cg07306253; while one hypomethylated CpG site (cg04977528) also has a protective effect. In contrast, one hypermethylated CpG site (cg26669793) and two hypomethylated CpG sites (cg25903363 and cg26628907) show damaging effects (Fig. 4b).

Contradicting the popular notion that most age-related differential methylation features are bad for the organism, our findings revealed that, in terms of the number of CpGs, there was no enrichment for either protective or damaging differential methylation during aging (Extended Data Fig. 8). Note that the age-associated CpG sites are identified in cross-sectional studies, therefore, a fraction of protective sites we observed could be explained by survival bias (i.e., CpG sites that promote late-life survival). Interestingly, there is a stronger depletion of meSNP in adaptive sites compared to the damaging sites, consistent with the notion that the adaptative mechanism is stronger regulated compared to the damage (Extended Data Fig. 2) ^30^. We also found that there was no significant correlation between the size of the causal effect and the magnitude of age-associated differential methylations (Fig. 4b, Extended Data Fig. 8), suggesting that CpG sites with a greater effect on healthy longevity do not necessarily change their level of methylation during aging. This result is consistent with our findings discussed above and explains the lack of enrichment of causal sites in existing epigenetic clocks.

The product of the causal effect and age-associated differential methylation (*b*_*age*_ × *b*_*MR*_) provides an estimate of the effect of age-related differential methylation on aging-related phenotypes in a unit of time. We calculated the cumulative effect of age-associated differential methylations on Aging-GIP1 by cumulative summing the effect of top 3,000 age-associated CpG sites, and calculated the empirical P-value through 10,000 permutations (Fig. 4c). Importantly, we discovered that although the number of protective and damaging CpG sites was similar, the cumulative effect of combined age-related DNA differential methylation is significantly detrimental to age-related phenotypes (P = 0.007), consistent with the overall damaging nature of aging.

### Developing causality-enriched epigenetic clocks

Although various existing epigenetic aging clock models can accurately predict the age of biological samples, they are purely based on correlation. This means that the reliability of existing clock models is highly dependent on the correlation structure of DNA methylation and phenotypes. This may result in unreliable estimates when extrapolating the model to predict the age of novel biological conditions (i.e., applying clocks to interventions that do not exist in the training population), as the correlation structure may be corrupted by the new intervention.

To overcome this problem, we developed novel epigenetic clocks that are based on putative causal CpG sites identified by EWMR (Fig. 5a). Specifically, we trained an elastic net model predicting chronological age on whole blood methylation data from 2,664 individuals ^31^, using CpG sites identified as causal to adjusted Aging-GIP1 by EWMR (adjusted *P* < 0.05). In regular epigenetic clock models, the penalty weight is defined to be 1 for all CpG sites, which produces models that are purely based on correlation. Instead, we introduced a novel causality-enriched elastic net model, where we assigned the feature-specific penalty factor based on the causality score for each CpG site (Method). The influence of the causality score on the feature-specific penalty factor is controlled by the causality factor τ, which is an adjustable parameter. We also constructed the model in the way that the CpGs that promotes healthy longevity are favored (i.e., with *β* > 0), to avoid including sites that disrupt the system but unrelated to aging. If τ = 0, the whole model is reduced to a regular elastic net regression, where the penalty factor equals one for all features. When τ becomes large, the model is more influenced by the causality score and tends to assign larger coefficients to the features with a higher causality score (Fig. 5A, Method).

**Fig. 5.**
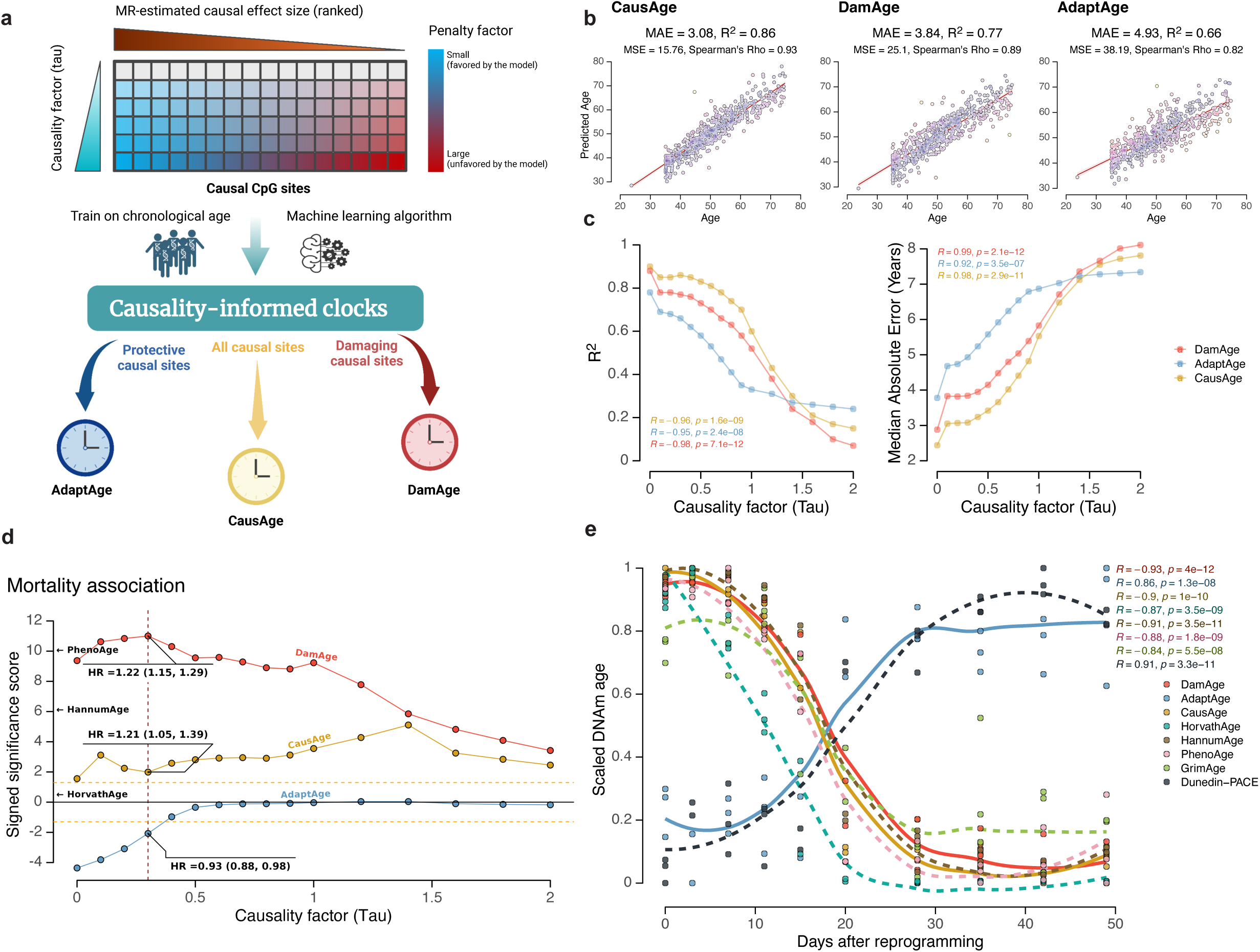
Construction and application of causality-enriched epigenetic clocks. **a.** Schematic diagram shows the procedure of constructing causality-enriched epigenetic clocks. **b.** Scatter plots show the accuracy of causal clocks on the test set. The X-axis shows the real age of each sample, and the Y-axis shows the predicted age of the same sample based on each clock model. Median absolute error (MAE) and Pearson’s R are annotated. **c.** Line plot showing the relationship between causality factor (τ) and clock accuracy measured by MAE and Pearson’s R. **d.** Line plot shows the relationship between the causality factor (τ) and -log10(p) for the association with mortality risk (signed by log2(hazard ratio)) estimated from the meta-analysis of FHS and NAS cohorts. Yellow dashed line shows the *P* threshold of 0.05. Hazard ratio of mortality risk for every 10-year increase in age for each clock model and the 95% confidential interval for τ = 0.3 is annotated. Results based on Horvath age, Hannum age, and PhenoAge are also shown by arrows for comparison. **e.** Scatter plots show the application of causal clocks and five other aging clocks to reprogramming of fibroblasts to iPSCs. X-axis shows days after initiating reprogramming. Pearson’s R and *P* values are annotated. All P-values reported are raw P-values without multiple testing correction, as the purpose is to compare across different clocks instead of deciding whether the test is significant.

Using this method, we trained the model to build the causality-enriched epigenetic clock CausAge (586 sites) using 2,664 blood samples. To separately measure adaptive and damaging DNA differential methylation during aging, we further separated putative causal CpG sites into two groups based on the causal effect size from MR and the direction of age-associated differential methylation (Fig. 4b). We then built DamAge, the damaging clock, which contains only the damaging CpG sites (1090 sites), and AdaptAge, the protective clock, which contains only the adaptive/protective CpG sites (1000 sites, Fig. 5a). We show that the model’s accuracy significantly decreased as the causality factor τ increased (Fig. 5b, c, Extended Data Fig. 9). This is because the causality factor τ controls the trade-off between the correlation and causality score-weighted penalty factor, and the causality score is not always correlated with the predictive power of age. For example, a CpG site with a high correlation with age may not be causal to aging, and *vice versa*. We therefore selected causality factor τ of 0.3 in the downstream analysis, which is the largest τ value with MAE < 5 years in the validation set and maximized the association with mortality (Fig. 5c-d).

### DamAge and AdaptAge uncouple damage and adaptation in aging

By design, AdaptAge contains only the CpG sites that capture protective effects against aging. Therefore, in theory, the subject predicted to be older by AdaptAge may be expected to accumulate more protective changes during aging. On the contrary, DamAge contains only the CpG sites that exhibit damaging effects, which may be considered as a biomarker of age-related damage. Therefore, we hypothesized that DamAge acceleration may be harmful and shorten life expectancy, whereas AdaptAge acceleration would be protective or neutral, which may indicate healthy longevity.

To test this hypothesis, we first analyzed the associations between human mortality and epigenetic age acceleration quantified by causality-enriched clocks using 4,651 individuals from the Framingham Heart Study, FHS offspring cohort (n = 2,544 Caucasians, 54% females, Supplementary Information) and Normative ageing study (NAS, n = 2,280, Supplementary Information). Among the three causality-enriched clocks, DamAge acceleration showed the strongest positive association on mortality (P = 9.9e-12) and outperformed CausAge (P = 0.01), AdaptAge (P = 0.008), Horvath clock (P = 0.34), Hannum clock (P = 8.2e-7), and is comparable to PhenoAge (P = 9.2e-11, Fig. 5d). This finding supports the notion that age-related damage is the main contributor to the risk of mortality, and the solely damage-base clock is better than the mixture of both damage and adaptation. In contrast, AdaptAge acceleration showed a significant negative association with mortality, suggesting that protective adaptations during aging, measured by AdaptAge, are associated with longer lifespan. In addition, epigenetic age accelerations measured by DamAge and AdaptAge were near-independent (Pearson’s R = 0.14, Extended Data Fig. 9). These findings highlight the importance of separating adaptive and damaging age-associated differential methylation when building aging clock models.

Interestingly, although the clock accuracy monotonically decreased as the causality factor τ increased, the association between mortality and epigenetic age acceleration did not follow the same trend (Fig. 5d). Especially for DamAge, the mortality association increased as the τ increased and peaked when τ was around 0.3. Also, DamAge consistently outperformed CausAge in predicting mortality risk, even though CausAge was more accurate in age prediction (Fig. 5b-e), the association between CausAge and mortality may be weakened due to the inclusion of adaptive sites. This suggests that although the introduction of the causality score and separation of damaging CpGs may decrease the accuracy of the clock in terms of predicting chronological age, it improves the prediction of aging-related phenotypes.

Induced pluripotent stem cell (iPSC) reprogramming is one of the most robust rejuvenation models, which was shown to be able to strongly reverse the epigenetic age of cells ^32^. We applied the causality-enriched clock models to reprogramming of fibroblasts to iPSC ^33^. For comparison, we also included five published epigenetic models, namely Horvath Age, Hannum Age, PhenoAge, GrimAge and DunedinPACE. The Horvath and Hannum clock were trained on chronological age ^32,34^, PhenoAge was trained on the age-adjusted by health-related phenotypes ^35^, GrimAge was trained on mortality ^36^, and DunedinPACE was trained to predict the pace of aging ^37^. Consistent with Horvath clock, Hannum clock, PhenoAge, and GrimAge, DamAge revealed that epigenetic age decreased during iPSC reprogramming, but with a stronger negative correlation with the time of reprogramming and higher statistical significance (R = -0.93, P = 4e-12, Fig. 5f). This observation suggests that DamAge may better capture the damage-removal effect of iPSC reprogramming. On the contrary, AdaptAge increased significantly during the reprogramming process (R = 0.86, P = 1.3e-8), suggesting that protective age-associated differential methylation does not capture the rejuvenation effect and that in fact cells may acquire even more protective changes during iPSC reprogramming.

To further examine how DamAge and AdaptAge capture age-related damage and protective adaptations, respectively, we tested the performance of causality-enriched clocks using various datasets. We first examined several aging-related conditions, namely atherosclerosis, cancer, and hypertension (Fig. 6a). We analyzed the blood samples from clinical atherosclerosis patients (n = 8) and healthy donors (n = 8) in the LVAD study ^38^. All eight clocks tested showed that the atherosclerosis patients are significantly older than healthy controls (Fig. 6a). We also analyzed 70 prostate cancer cases with good or poor prognosis ^39^. Only DamAge successfully detected a significant age acceleration in patients with bad cancer prognosis (P = 0.039), while Hannum age detected a significant inverse effect where the patients with good prognosis were age accelerated (P = 0.044). For hypertensive heart disease, we analyzed blood samples from 44 hypertensive patients and 44 healthy controls ^40^. Both CausAge and DamAge showed significant age acceleration in hypertensive patients (CausAge P = 0.002, DamAge P = 0.04). Similar effects could be detected with GrimAge (P = 0.002) and DunedinPACE (P = 0.02), but not with AdaptAge, PhenoAge, and two 1st generation clocks. These results suggest that DamAge could more robustly represent the effect of age-related conditions, compared to the published 1st and 2nd generation clocks.

**Fig. 6.**
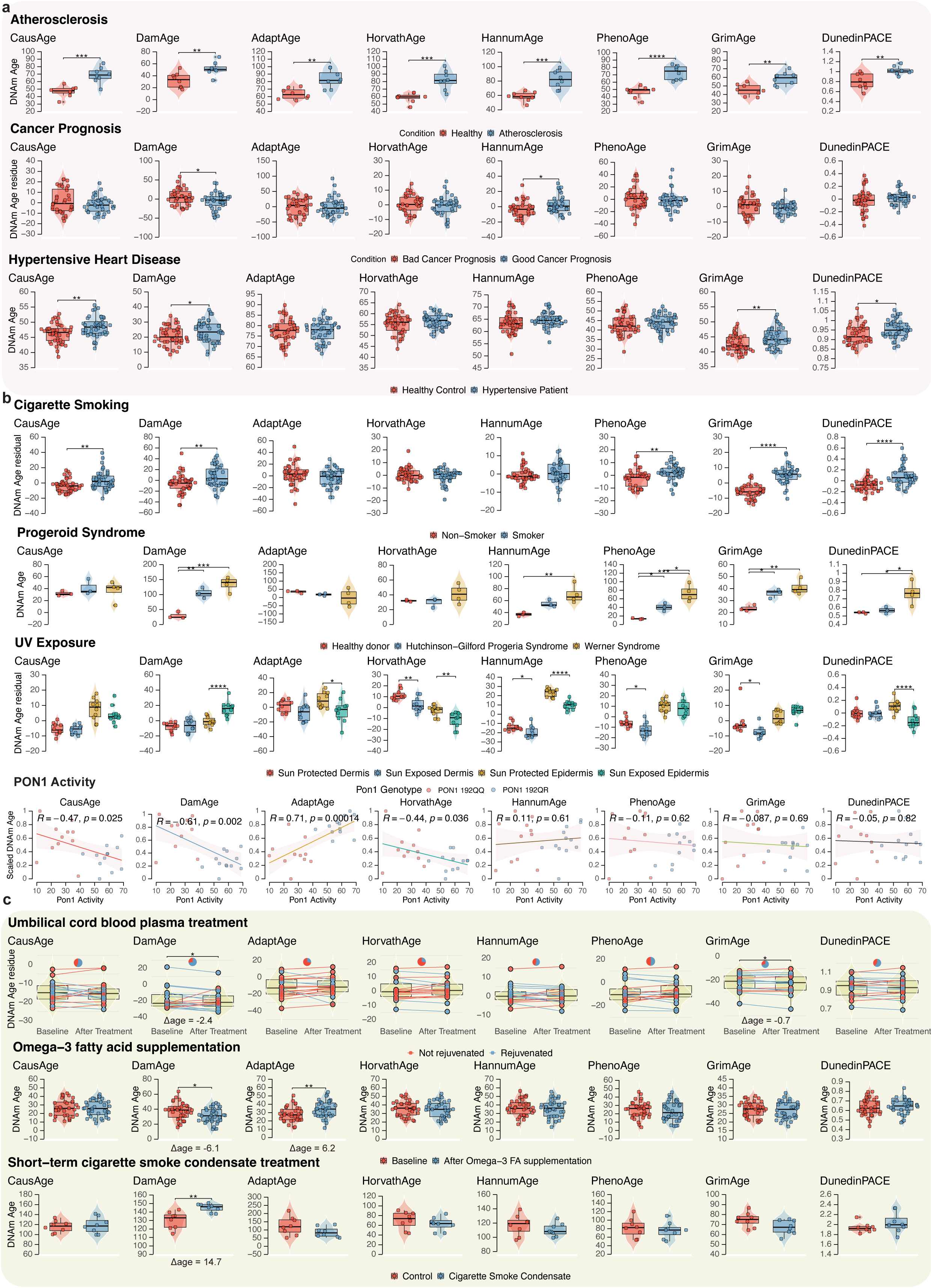
Causality-enriched epigenetic clocks can better capture aging-related effects. **a.** Box plots show the association between epigenetic age and aging-related conditions, including atherosclerosis (N = 16, significant P values from left to right are 2.79e-4, 0.00619, 0.0011, 2.77e-4, 2.81e-4, 2.75e-5, 0.00231, 0.00413), prostate cancer prognosis (N = 70, significant P values are 0.0394 and 0.044), and hypertensive heart disease (N = 88, significant P values are 0.002, 0.049, 0.002, 0.017). **b.** Box plots show the association between epigenetic age and damaging conditions, including smoking (n = 80, significant P values are 0.004, 0.006, 0.005, 1.01e-14, 7.16e-6), Progeroid Syndromes (n = 10, significant P values are 0.004, 5.55e-4, 0.014, 0.047, 0.002, 0.033, 0.028, 0.006, 0.033, 0.04), and Sun exposure (n = 40, significant P values are 6.1e-4, 7.4e-6, 8.48e-4, 1.18e-5, 7.66e-7, 3.96e-7, 9.78e-5, 0.0256, 2.39e-5, 0.00576, 1.38e-9, 2.82e-6, 4.99e-17, 7.4e-8, 8.91e-5, 1e-8, 1.24e-5, 7.55e-6, 0.0122, 9.47e-4, 1.46e-4). Scatter plots show the correlation between epigenetic age and blood PON1 activity (n = 48). Epigenetic age prediction is rescaled to a 0-1 scale for better comparison. The color scheme shows the PON1 genotype in subjects. Linear regression is performed, and Pearson’s R and P values are annotated. **c.** Box plots show the association between epigenetic age and short-term treatments, including the umbilical cord blood plasma treatment (n = 18, significant P values are 0.0481 and 0.0481), 15 months of cigarette smoke condensate (CSC) treatment (n = 16, significant P value is 0.002), and 6-week supplementation of overweight subjects with omega-3 fatty acids (n = 69, significant P value is 0.0087).The effect size of significant effect is annotated. For umbilical cord blood plasma treatment, one-tailed paired sign-test was performed, and the color scheme and the pie chart indicate whether the subject is rejuvenated after treatment based on the corresponding clock. For unpaired box plots, significant pairs based on two-tailed t-test are annotated with stars. * *P* < 0.05, ** *P* < 0.01, *** *P* < 0.001, **** *P* < 0.0001. All reported P values are corrected for multiple testing using FDR method within each clock group. Box plots indicate the median (central line), 25th and 75th percentiles (bounds of box), and the minimum and maximum values (whiskers).

Next, we examined conditions that specifically promote age-related damage (Fig. 6b). Smoking is a well-known risk factor for many age-related diseases, and it also causes DNA damage and oxidative stress. We compared the epigenetic age of smokers (n = 40) and non-smokers (n = 40) ^41^. CausAge (P = 0.004) and DamAge (P = 0.006), together with all three 2nd generation clocks could detect significant age-acceleration among smokers, while AdaptAge and two 1st generation clocks did not. Progeroid syndrome is a group of rare genetic disorders that cause premature aging. We analyzed blood cell samples from healthy donors (n = 3), and patients with Hutchinson-Gilford Progeria Syndrome (HGP, n = 3) and Werner Syndrome (n = 4) ^42^. We observed significant DamAge acceleration in both HGP (P = 0.004) and Werner Syndrome (P = 5e-4) compared to healthy controls. Similar effects were detected also with PhenoAge and GrimAge. Hannum age and DunedinPACE detected age acceleration in Werner Syndrome but not in HGP, while no significant effect was found by other clocks (Fig. 6b). We then analyzed dermis and epidermis samples with or without sun exposure (n = 10 per group) in old adults (age > 60) ^43^. As the exposure to ultraviolet promotes DNA damage and aging, it may be considered a model of age-related damage. As expected, we observed significant DamAge acceleration in sun-exposed epidermis compared to sun-protected epidermis (P = 2e-5), while no significant effect was observed in the dermis tissue. AdaptAge of the sun-exposed epidermis was significantly lower (P = 0.01). Surprisingly, based on most other published clocks (including Horvath age, Hannum age, and DunedinPACE), the sun-exposed epidermis was predicted to be significantly younger than sun-protected epidermis. Only GrimAge showed the expected effect direction but did not reach statistical significance (P = 0.1).

Paraoxonase 1 (PON1) is one of most studied genes associated with cardiovascular disease, oxidative stress, inflammation, and healthy aging. Specifically, PON1 plays an important role in detoxifying organophosphorus compounds and removing harmful oxidized lipids ^44^. The genetic variant of PON1 (R192Q) significantly decreases PON1 activity and is known to be associated with an increased risk of cardiovascular disease and neurodegenerative diseases ^45^. Interestingly, the PON1 Q allele is significantly depleted in centenarians ^45^. We analyzed the relationship between PON1 activity and epigenetic age in 48 whole blood samples (Fig. 6a) ^46^. DamAge shows a significant negative correlation with PON1 activity (R = -0.55, p = 0.0062), whereas AdaptAge showed a significant positive correlation with PON1 activity (R = 0.69, p = 0.0003). Again, this association was not observed by other epigenetic clocks, except for Horvath age, but with a less significant negative correlation (P = 0.04). Together, we showed that DamAge can reliably detect damage-related biological age acceleration.

Causality-enriched clocks could also capture the aging-related effects of short-term interventions. We first examined the effect of human umbilical cord plasma concentrate injection, which was reported to have age reversal effects ^47^. In this study, 18 older participants were treated with human umbilical cord plasma concentrate injection weekly (1 ml intramuscular) over a 10-week period. We found that this rejuvenation effect could only be captured with DamAge (P = 0.04) and GrimAge (P = 0.04, Fig. 6c). Note that this effect is not detected by other clocks, including AdaptAge, which may be contributed by the weak effect and short treating interval. Similarly, a 6-week omega-3 fatty acid supplementation in overweight subjects (n = 34) ^48^, which was shown to be protective against age-related cardiovascular diseases, significantly increased AdaptAge (P = 0.009) and reduced DamAge (P = 0.02, Fig. 6c). We also found that short-term treatment with cigarette smoke condensate in bronchial epithelial cells significantly accelerated DamAge (P = 0.002) but did not affect other tested clocks (Fig. 6c). Together, our data demonstrate the importance of separating damage and adaptation when building biomarkers of aging and provide novel tools to quantify aging and rejuvenation.

Small for gestational age (SGA) is a condition defined as birth weight less than the 10th percentile for gestational age, usually due to limited access to the nutrient. We found that children with SGA have a significantly lower DamAge and higher AdaptAge than children with normal birth weight (GSE47880, Extended Data Fig. 10). These effects were not captured by other epigenetic clocks tested. SGA is usually considered a pathological condition; some studies suggest that it may be because early life benefits can be reversed in later life by exposure to excess nutrients. The different roles of SGA in the early and late stages of life may need to be further investigated in future studies.

In vitro fertilization (IVF) is a common method of treating infertility. Yet, previous studies have shown that IVF may increase the risk of perinatal morbidity and mortality. We analyzed the DNA methylation data from neonatal blood spots of 137 newborns conceived unassisted (NAT), through intrauterine insemination (IUI), or through IVF using fresh or cryopreserved (frozen) embryo transfer ^49^. We found that IVF-conceived newborns using fresh or cryopreserved embryos had higher DamAge acceleration and lower AdaptAge than NAT-conceived newborns (Extended Data Fig. 10). On the other hand, IUI-conceived newborns showed no differences in their DamAge and AdaptAge with controls (Extended Data Fig. 10). This effect could not be observed by other five epigenetic clocks tested, except for Horvath age. We also analyzed peripheral blood DNA methylation data from patients with single-locus or Multi-loci imprinting disturbances (SLID or MLID), which is the condition of losing methylation at single or multiple imprinting centers ^50^. Similar to IVF, we found that patients with imprinting disorders showed significantly higher DamAge and lower AdaptAge (Extended Data Fig. 10). Together, these results suggest that DamAge and AdaptAge may serve as preferred biomarkers for the events affecting aging traits during development.

## Discussion

Epigenetic aging clocks, like those based on DNA methylation patterns, are adept at predicting the age of biological samples ^8^. DNA methylation changes with age, influencing chromatin structure and gene expression, potentially affecting aging phenotypes. For instance, DNA methylation may play a causal role in cellular rejuvenation during iPSC reprogramming ^32^. However, it’s crucial to discern if age-related differential DNA methylation causes aging phenotypes or merely the downstream effect of them. Transcriptome-wide Mendelian Randomization (MR) studies suggest that differentially expressed genes in diseases often result from, rather than cause, the disease ^28^. Our epigenome-wide MR (EWMR) supports this, showing minimal overlap between CpG sites causally linked to healthy longevity and those changing with age.

MR is a powerful method to identify causal relationships between exposure traits and phenotypes ^14^. The causality between the exposure and outcome in MR studies relies on three key assumptions: First, the genetic variants used as instrumental variables (IVs) must be robustly associated with the exposure of interest. Second, the IVs should not be associated with any confounding factors that influence both the exposure and the outcome. Third, the association between the IVs and the outcome must be mediated solely through the exposure, not through any alternative pathways. In our study, we used SNPs associated with DNA methylation levels at specific CpG sites as IVs to investigate their causal relationship with aging-related outcomes. While MR provides a framework to infer causality, it is crucial to recognize its limitations, particularly in the context of complex traits like aging. Our approach, although comprehensive, does not definitively establish causality but rather suggests putative causation based on the MR framework. The assumption that the effect of SNPs on aging-related outcomes is mediated exclusively through methylation changes at CpG sites is challenging to validate and may not encompass all underlying mechanisms. Hence, while our results contribute to the understanding of the relationship between DNA methylation and aging, they should be interpreted with caution, recognizing inherent limitations of the MR approach in fully elucidating causal pathways in the complex landscape of aging biology.

MR is also limited by the availability of genetic instruments for those traits ^14^. In our study, we used DNA methylation quantitative trait loci (meQTLs) from the Illumina 450K array as instruments. But, the high correlation between nearby CpG sites and the scarcity of mutations at causal CpG sites (meSNPs) make it challenging to separate individual CpG effects. Hence, identified CpG sites are more likely markers for causal regions in aging-related phenotypes. While our findings mainly pertain to blood, as our genetic instruments were derived from the largest meQTL study in blood (GoDMC), a significant proportion of cis-meQTLs are tissue-conserved, suggesting broader relevance ^50^.

Considering molecular changes in aging for intervention targets may be flawed, as correlation doesn’t imply causation. Many molecular changes could be neutral or adaptive responses to aging. For example, declines in IGF-1 and growth hormone, and increased protein aggregation in C. elegans, are adaptive mechanisms extending lifespan ^43,51^. Our results indicate such adaptations at the epigenetic level are as common as damaging changes. Hence, focusing on causal effects and their direction is crucial.

Our causal epigenetic clock models (CausAge, AdaptAge, DamAge) distinguish protective changes from damaging ones. AdaptAge, for instance, predicts lower mortality risk in individuals with elevated protective adaptations. Interestingly, DamAge and AdaptAge, while accurately predicting chronological age, reflect different biological meanings in their delta-age terms. Their correlation in the general population may stem from survival and collider biases, highlighting the need for careful interpretation of epigenetic age acceleration in clinical settings.

In conclusion, our causality-enriched clock models offer new insights into aging mechanisms and the evaluation of interventions. They underscore the importance of distinguishing between adaptive and damaging epigenetic changes in aging research. Future studies, especially in diverse tissues, are needed to expand and validate these findings.

## Methods

The research conducted in this study adheres to all applicable ethical regulations. This study, due to its nature, did not require ethical review or approval by an institutional review board.

### Epigenome-wide Mendelian Randomization analysis

In MR analysis, the definition of causal relationship is that associations of SNPs with CpG methylation are directionally consistent and proportional in magnitude to associations of SNPs with aging-related phenotypes. Genetic variants that are strongly associated with whole blood DNA methylation level (FDR < 0.05) were used for the MR analysis. Only meQTLs in the cis-acting regions were used to avoid pleiotropic effects. As the generalized MR method achieves a higher statistical power by including partially correlated instruments while accounting for the LD structure, we used LD clumping to only remove meQTLs with strong LD (r^2^ > 0.3), as suggested by Burgess et al. ^52^. Three MR methods were used for the main analyses: Wald ratio when only one meQTL was available, generalized inverse variance weighted (gIVW) when at least two meQTLs were available, and generalized MR-Egger regression (gEgger) when at least three meQTLs were available ^52^. The MR analyses were conducted using the MendelianRandomization R package and TwoSampleMR R package ^53,54^.

We only included cis-meQTLs (meQTLs located within 2 MB of target CpG sites) in our analysis to avoid pleiotropic effects, as they are more likely to affect DNA methylation via direct mechanisms. To remove additional pleiotropic effects, we used the results of gEgger, whose estimate is robust to directional pleiotropic effects if the significant intercept is detected by gEgger regression (*P* < 0.05).

CpG-phenotype pairs with *P*_adjusted_ < 0.05 after Bonferroni correction were used to select causal CpG sites with the strongest MR evidence. All CpG-phenotype pairs with FDR < 0.05 were considered potential causal CpG sites and used in the downstream sensitivity analysis.

### Mediation analysis

We conducted multivariable MR (MVMR) to dissect significant CpG-phenotype causal effects (θ_T_) into direct effects (θ_D_) and indirect effects through transcript levels following the methodology outlined in Sadler et al., 2022 and using the smr-ivw software (https://github.com/masad-ler/smrivw)^26^. Genetic effect sizes on CpGs (mQTLs) came from the GoDMC consortium (N=32,851) ^12^, and on transcript levels (eQTLs) from the eQTLGen consortium (N=τ31,684) ^55^, both derived from whole blood. Mediation analyses were assessed for CpG-Aging-GIP1 and CpG-adjusted Aging-GIP1 pairs.

Transcript mediators were selected to be in *cis* (< 500kb away from the CpG site) and causally associated to the CpG. This latter condition was satisfied when univariable MR effects from the CpG site on the transcript had an MR p-value < 0.01. Instrumental variants were required to be associated to either the CpG or included transcripts and as in the univariable MR analysis they were selected to be correlated at r^2^ ≤ 0.3. The mediation proportion (MP) was estimated as 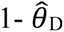/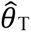.

### Causality-enriched epigenetic clock model

Elastic net regression is a regularized linear model that solves the problem.

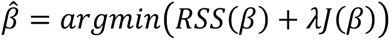

where *RSS*(*β*) is the residual sum of squares, λ is the regularization parameter, and *J*(*β*) is the regularization term ^56^. The *J*(*β*) term is the sum of the L1 and L2 terms, which is defined as

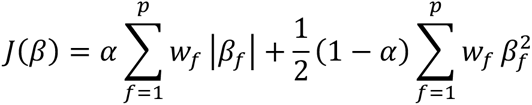

The parameter is the elastic net mixing parameter, which controls the balance between the L1 and L2 terms. *w*_*f*_ is the penalty factor for each feature we introduced. In a regular epigenetic clock model, *w*_*f*_ is defined to be 1 for all the features, which produces the model that is purely based on correlation. We introduced a causality-enriched elastic net model, where we defined the feature-specific penalty factor as

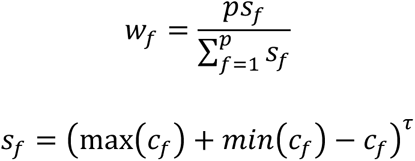

Here the *c*_*f*_ is the absolute value of the causality score for each feature, which is calculated from the causal effect size from MR weighted by colocalization probability. The τ > 0 is a tuneable parameter that controls how much the causality score affects the feature-specific penalty factor. If τ = 0, the whole model is reduced to a regular elastic net regression, where *w*_*f*_ = 1 for all the features. When τ becomes large, the model is more influenced by the causality score and tends to assign larger coefficients to the features with a higher causality score. To balance the precision and causality, we defined τ as 0.3, which is the largest τ value with MAE < 5 years in the validation set and maximized the association with mortality. In addition, we introduced a hyperparameter array to control the limits of each model weight, ensuring the range and sign of the weight align with the patterns of aging. The upper limits for the sites that decrease healthy longevity are defined to be a small value (% of the site by default) to encourage the models rely more on CpG sites that promotes healthy longevity.

Using this method, we trained the model on whole-blood methylation data from 2,664 individuals ^57^. We built the causality-enriched epigenetic clock model CausAge. To separately measure adaptive and damaging DNA methylation changes during aging, we further separate the causal CpG sites into two groups based on causal effect size from MR and the direction of age-related changes. We then built DamAge, a damaging clock, and AdaptAge, a protective clock.

### Genetic correlation analysis

Genetic correlation between traits related to aging is calculated using the LD score regression (LDSC) ^58^. SNPs that were imperfectly imputed with INFO less than 0.9 or with a low minor allele frequency less than 5% were removed to reduce statistical noise. LDSC was performed using LDSC software v1.0.1 (https://github.com/bulik/ldsc).

### Sensitivity analyses

*Horizontal pleiotropy*. The horizontal pleiotropic effect in instrumental variants may cause biased causal effect estimation from the gIVW method. To detect unbalanced horizontal pleiotropy among genetic instruments, we used the intercept gEgger regression, which provides an estimate of the directional pleiotropic effect ^59^. Note that by including a partially correlated instrument, the gEgger intercept also has more statistical power to detect the pleiotropic effect. CpG-phenotype pairs with gEgger intercept *P* value less than 0.05 were potentially affected by the pleiotropic effect, and the gIVW method may be biased. We, therefore, reported an estimate and *P* value from the gEgger method instead of the gIVW method for these MR signals, as the gEgger estimate is robust to the horizontal pleiotropic effect ^59^.

### Heterogeneity

To detect heterogeneity of the MR estimates in each meQTL, we performed the Cochran’s Q test and the Rücker’s Q test for the gIVW and gEgger results, respectively. Since heterogeneity does not necessarily affect causal effect estimation, we kept the MR signals heterogeneous while reporting potential heterogeneity in the result table.

### Directionality test

To exclude MR signals caused by reverse causality (i.e., methylation changes caused by outcome phenotype), we applied MR Steiger test ^21^, which is the method to test the directionality of causal effect estimated by MR. We then removed all MR signals with reverse directionality. We also applied the cis-reverse MR on the CpG-outcome pairs if the genome-wide significant instrument for the outcome traits is available. The CpG-outcome pairs with correct causal effect direction and non-significant reverse causal effects are marked as with correct causal direction.

### Colocalization analysis

All MR signals that passed the FDR threshold of 0.05 were then subjected to the colocalization analysis. We applied pairwise a conditional and colocalization (PWCoCo) analysis ^60^, which is a powerful genetic colocalization approach that is able to detect multiple independent genetic signals. We considered colocalization probability (PP.H4) > 70% as strong evidence of colocalization. Also, since aging-related GWAS are in general noisy while cis-meQTL usually have strong genetic signals, colocalization probability tends to be low, and the probability of only having a signal from meQTLs (PP.H1) tends to be high. To overcome bias due to imbalanced power between exposure and outcome traits, we considered a conditional colocalization probability (conditional PP.H4 = PP.H4 / PP.H3+H4) by assuming that the aging-related trait always has genetic signals in the region when a significant MR signal is detected. We then also reported CpG-phenotype pairs with conditional PP.H4 > 70% as a potentially colocalized signal.

### Mortality association analysis

To evaluate our novel causal clocks for predicting all-cause mortality risk, we applied the clocks to a large-scale dataset comprising 4,651 individuals from (a) the Framingham Heart Study, FHS offspring cohort (n = 2,544 Caucasians, 54% females) ^15,16^ and (b) Normative ageing study (dbGaP accession phs000853.v2.p2)^61^. Methylation levels were profiled in blood samples based on Illumina 450k arrays. In FHS, the mean (SD) chronological age at the time of the blood draw was 66.3 (8.9) years old. During follow-up, 330 individuals died. The mean (SD) follow-up time (used for assessing time-to-death due to all-cause mortality) was 7.8 (1.7) years. As noted, this AgeAccel measure is independent of chronological age. Next, we applied Cox regression analysis for time-to-death (as a dependent variable) to assess the predictive ability of our causal clocks for all-cause mortality, using the AgeAccel measures. The analysis was adjusted for age at the blood draw and adjusted for gender and batch effect in FHS. We combined the results across FHS and NAS cohorts by fixed effect models weighted by case number. The meta-analysis was performed in the R metafor function.

### Statistics & Reproducibility

The Methods and Extended Methods sections outline statistical approaches employed in this study for data analysis. Data processing and analysis were primarily performed using R version 4.0.2, except where noted. In t-tests performed, data distribution was assumed to be normal but this was not formally tested. Given the nature of this study, predetermining sample sizes, randomizing experiments, and blinding investigators to experimental conditions were not relevant or applicable. In the clock training step, the CpG sites with less than 90% coverage were excluded from the analysis during quality control.

## Data Availability

All analyses in this study were conducted using publicly available data. The datasets used are listed in Table 1 and their respective URLs are: Longevity GWAS summary statistics: https://www.lon-gevitygenomics.org/downloads. Parental lifespan GWAS summary statistics: https://datashare.ed.ac.uk/handle/10283/3209. Healthspan GWAS summary statistics: https://www.gwasarchive.org/. Frailty index GWAS summary statistics: https://figshare.com/arti-cles/dataset/Genome-Wide_Association_Study_of_the_Frailty_Index_-_At-kins_et_al_2019/9204998. Epigenetic age acceleration GWAS summary statistics: https://datashare.ed.ac.uk/handle/10283/3645. GEO datasets GSE107143, GSE127985, GSE192918, GSE193795, GSE210245, GSE51954, GSE94876, GSE98056, GSE101673, GSE78773, GSE90117, GSE79257, GSE42865. Any other data generated in this study upon which conclusions are based are available in Supplementary Tables.

## Code Availability

Mendelian Randomization analyses were conducted using the R package TwoSampleMR version 0.5.6: https://mrcieu.github.io/TwoSampleMR/; R package MendelianRandomization version 0.7.0: https://cran.r-project.org/web/packages/MendelianRandomization/index.html; Genetic cor-relation analysis was performed using: LDSC software v1.0.1: https://github.com/bulik/ldsc. Mediation analysis was performed using smr-ivw v1.0 https://github.com/masadler/smrivw. Colocalization analysis was performed using PWCoCo v1.0 https://github.com/jwr-git/pwcoco. Elastic net model was trained using glmnet v4.1 https://cran.r-project.org/web/packages/glmnet/in-dex.html. Custom code used are available in Supplementary Information. Algorithm of CausAge, DamAge, and AdaptAge are available in Supplementary information, as well as ClockBase (www.clockbase.org) and bio-learn python package (https://bio-learn.github.io/) ^62,63^.

## Acknowledgments

We thank the DNA Methylation Consortium (GoDMC) for releasing the summary statistics of meQTLs. We also thank Csaba Kerepesi, Marco Mariotti, Daniel L. McCartney, and Riccardo E. Marioni for their help and advice during the initial stages of this study. We specially thank Yin Fang for the artwork design. This study is supported by the National Institute on Aging, Impetus grants and the Michael Antonov Foundation. The Framingham Heart Study is funded by National Institutes of Health contract N01-HC-25195 and HHSN268201500001I. The US Department of Veterans Affairs (VA) Normative Aging Study is supported by the National Institute of Environmental Health Sciences (NIEHS) (R01ES015172, R01ES021733) as well as by the Cooperative Studies Program/ERIC, US Department of Veterans Affairs, and is a research component of the Massachusetts Veterans Epidemiology Research and Information Center (MAVERIC). Additional support to the VA Normative Aging Study was provided by the US Department of Agriculture, Agricultural Research Service (contract 53-K06-510).

## Author Contributions

K.Y. initiated the study and performed data collection and analyses. V.N.G. supervised this research and provided funding. H.L., M.C.S., and A.T. were involved in data analysis. A.T.L., M.M., and S.H. contributed to data interpretation. Z.K. and X.S. assisted in methodology refinement and statistical analysis. All authors contributed to writing and revising the manuscript and approved the final version for publication.

